# Engineered chimeric T cell receptor fusion construct (TRuC)-expressing T cells prevent translational shutdown in SARS-CoV-2-infected cells

**DOI:** 10.1101/2021.06.25.449871

**Authors:** Ira Godbole, Kevin Ciminski, O. Sascha Yousefi, Salma Pathan-Chhatbar, Deniz Saltukoglu, Niklas Vesper, Pavel Salavei, Juliane Strietz, Nicole Gensch, Michael Reth, Martin Schwemmle, Wolfgang W. Schamel

## Abstract

SARS-CoV-2, the causative agent of Covid-19, is known to evade the immune system by several mechanisms. This includes the shutdown of the host cellular protein synthesis, which abrogates the induction of antiviral interferon responses. The virus initiates the infection of susceptible cells by binding with its spike protein (S) to the host angiotensin-converting enzyme 2 (ACE2). Here we applied the T cell receptor fusion construct (TRuC) technology to engineer T cells against such infected cells. In our TRuCs an S-binding domain is fused to the CD3ε component of the T cell receptor (TCR) complex, enabling recognition of S-containing cells in an HLA independent manner. This domain either consists of the S-binding part of ACE2 or a single-chain variable fragment of an anti-S antibody. We show that the TRuC T cells are activated by and kill cells that express S of SARS-CoV-2 and its alpha (B.1.1.7) and beta (B.1.351) variants at the cell surface. Treatment of SARS-CoV-2 infected cells with our engineered T cells did not lead to massive cytotoxicity towards the infected cells, but resulted in a complete rescue of the translational shutdown despite ongoing viral replication. Our data show that engineered TRuC T cell products might be used against SARS-CoV-2 by exposing infected cells to the host innate immune system.

## Introduction

Covid-19 is caused by a new virus of the beta-coronavirus family, the severe acute respiratory syndrome coronavirus 2 (SARS-CoV-2). Most infections are rather mild and are controlled by adaptive immune responses. Both, strong CD4 and CD8 responses as well as neutralizing antibodies are important (Grifoni et al., 2020; Schulien et al., 2021; Zheng et al., 2020) and might provide a certain degree of protection for re-infections. However, a significant proportion of patients require hospitalization and mortalities occur, especially in patients with pre-existing medical conditions, such as obesity, cardiovascular disease or type 2 diabetes, immunocompromised individuals (Docherty et al., 2020; Richardson et al., 2020) or those of older age (Docherty et al., 2020; Grasselli et al., 2020). In several severe cases SARS-CoV-2-specific CD4+ and CD8+ T cell responses failed to develop although neutralizing antibodies were found, but were insufficient to control disease (Rydyznski Moderbacher et al., 2020). Indeed, cases were reported in which T cells could control the infection well in the absence of detectable antibodies (Rydyznski Moderbacher et al., 2020). With more than 3.7 million deaths word-wide (June 2021) there is an urgent need for the treatment of ongoing severe COVID-19 cases, and personalized treatments might be required.

To enter host cells, the receptor binding domain (RBD) of the SARS-CoV-2 spike protein (S) binds to the host cell angiotensin-converting enzyme 2 (ACE2) (Wrapp et al., 2020; Yan et al., 2020; Zhou et al., 2020). Cleavage of S by the cellular serine protease TMPRSS2 at the cell surface initiates membrane fusion, so that the viral RNA is released into the cytosol for viral replication (Hoffmann et al., 2020). Alternatively, in the absence of TMPRSS2, the virion is endocytosed and S cleavage as well as membrane fusion takes place in endosomes (Hoffmann et al., 2020; Simmons et al., 2005).

One of the major innate antiviral defence mechanisms is the type I interferon (IFN-I) system, the induction of which stimulates the expression of a large number of interferon-stimulated genes (ISGs). In case of coronaviruses the viral RNA is sensed by retinoic acid-inducible gene I (RIG-I), thereby activating innate immune signaling (Hu et al., 2017; Sparrer and Gack, 2015). However, these viruses evade the innate immune response by abrogating host mRNA translation, thus preventing expression of IFN-I and ISGs (Hsu et al., 2021; Kamitani et al., 2009; Lei et al., 2020; Narayanan et al., 2008; Puray-Chavez et al., 2020; Thoms et al., 2020; Xia H et al., 2020; Yuen et al., 2020). In SARS-CoV-1 and SARS-CoV-2 multiple viral proteins are responsible for this translational shutdown (Hsu et al., 2021; Kamitani et al., 2009; Narayanan et al., 2008; Puray-Chavez et al., 2020; Thoms et al., 2020). For example, the non-structural protein 1 (Nsp1) inhibits nuclear export of host mRNA, but also blocks ribosomal translation by binding to the 40S ribosomal subunit (Kamitani et al., 2009; Narayanan et al., 2008; Thoms et al., 2020; Zhang et al., 2021) (Lapointe et al., 2021; Yuan et al., 2020b). In addition, Nsp5, Nsp14 and Nsp15 can also halt translation of host mRNA (Hsu et al., 2021; Lei et al., 2020; Xia H et al., 2020; Yuen et al., 2020). Indeed, SARS-CoV-2 infection as well as overexpression of these Nsps in Vero E6 or HEK293T cells inhibits the production of IFN-I and induction of ISGs (Lei et al., 2020; Puray-Chavez et al., 2020; Thoms et al., 2020; Xia H et al., 2020; Yuen et al., 2020). However, viral proteins can still be produced, most likely due to a special feature of the 5’ untranslated region of viral mRNAs (Tanaka et al., 2012). Thus, impairment of host mRNA translation may allow SARS-CoV-2 to evade host immunity. How host cells can counteract this translational shutdown is poorly understood.

T cells play an important role in the protection from infections and tumors through their production of cytokines and other immune mediators as well as through their cytotoxic activity. However, in chronic or severe viral infections as well as in the development of tumors, T cells have failed to control the disease. Thus, one possible approach for a treatment is to strengthen the T cell response by engineering the T cells to better recognize the target cells (infected or tumor cells) and to mount a stronger, yet more balanced immune response.

One strategy is to redirect polyclonal T cells to recognize tumor cells by expressing a chimeric antigen receptor (CAR) in the patients T cells. CARs consist of a tumor antigen-binding extracellular part, typically a single-chain variable fragment (scFv) of an antibody, a transmembrane domain and cytoplasmic regions derived from the TCR’s CD3ζ chain and co-stimulatory receptors, such as CD28 or 4-1BB (Guedan et al., 2019; Rafiq et al., 2020). Thus, the engineered T cells recognize tumors in an HLA-independent manner and couple this to T cell activation signals. CAR T cells are highly effective at eradicating certain hematological malignancies, such as acute B-lymphoblastic leukaemia and non-Hodgkin’s lymphoma, but fail against solid tumors (Glover et al., 2021). Further, over-activation and the dangerous cytokine release syndrome are important safety concerns (Fitzgerald et al., 2017; Morris et al., 2021).

A significant improvement of the CAR design was to employ the complete TCR machinery (Baeuerle et al., 2019; Hardy et al., 2020; Liu et al., 2021; Rana et al., 2021; Xu et al., 2018), instead of only the isolated cytoplasmic tail of CD3ζ. One such example are chimeric receptors called TCR fusion constructs (TRuCs). The TCR itself is a complicated multiprotein machine composed of the antigen-binding TCRαβ subunits and the signal transduction CD3εγ, CD3εδ and ζζ subunits. Ligand-binding leads to the exposure of signaling motifs in the cytoplasmic tails of CD3, such as a proline-rich sequence, basic-rich sequences, the receptor-kinase motif and the immuno-tyrosine based activation motifs (ITAMs) (Hartl et al., 2020; Reth, 1989; Schamel et al., 2019). These conformational changes, the recruitment of signaling molecules to the motifs and phosphorylation of the ITAMs are delicately controlled to determine the quantity and quality of the signal, eventually leading to the activation and differentiation of the T cell in a proper way. Importantly, mechanisms also control that in the absence of ligand the TCR is held in a closed, inactive conformation (Swamy, Schamel JI).

In TRuCs, the anti-tumoral scFv is directly engineered onto the TCR (Baeuerle et al., 2019; Gil et al., 2002; Minguet et al., 2007; Schamel and Reth, 2012). Thus, in stark contrast to the CARs, TRuCs use the full signaling potential of the TCR. In case of the εTRuC, where the scFv is fused to the ectodomain of the TCR’s subunit CD3ε, it was shown that the εTRuC T cells had a more physiological and controlled signaling than CAR T cells (Baeuerle et al., 2019; Rana et al., 2021) and were consequently more effective against B cell lymphoma and leukemia (Baeuerle et al., 2019). Importantly, the εTRuCT cell were also effective against solid tumors (Baeuerle et al., 2019; Liu et al., 2021) - situations where CAR-T cells fail.

We sought to apply the lessons learned from engineering anti-tumor εTRuC T cells to address viral infections, with SARS-CoV-2 as a model case. Hence, the aim of this study was to construct and test the ability of εTRuC T cells to control and counteract SARS-CoV-2 viral infection *in vitro.*

## Results

### SARS-CoV-2-specific ε TRuCs redirect T cells to recognize the S protein

To re-program T cells to recognize SARS-CoV-2-infected cells, we here constructed εTRuCs able to bind the SARS-CoV-2 S protein. To this end, the S-binding part of the extracellular domain of human ACE2 (aa 18-615) (Yan et al., 2020) was genetically fused to human CD3ε in a lentiviral vector. A short or a long linker were used to connect the two proteins, resulting in ACE2s-εTRuC and ACE2l-εTRuC (Fig. 1A and B). Besides ACE2, the human antibody CR3022 also binds to SARS-CoV-2 S (ter Meulen et al., 2006; Wrapp et al., 2020; Yuan et al., 2020a). Thus, a single chain fragment of the V_L_ and V_H_ regions (scFv) of CR3022 was fused to CD3ε, yielding the αS-εTRuC (Fig. 1A and B); the linker that was used before to construct the αCD19-εTRuC (Baeuerle et al., 2019) was chosen. All lentiviruses also encode for GFP fused with a T2A peptide to the εTRuC.

**Fig. 1.**
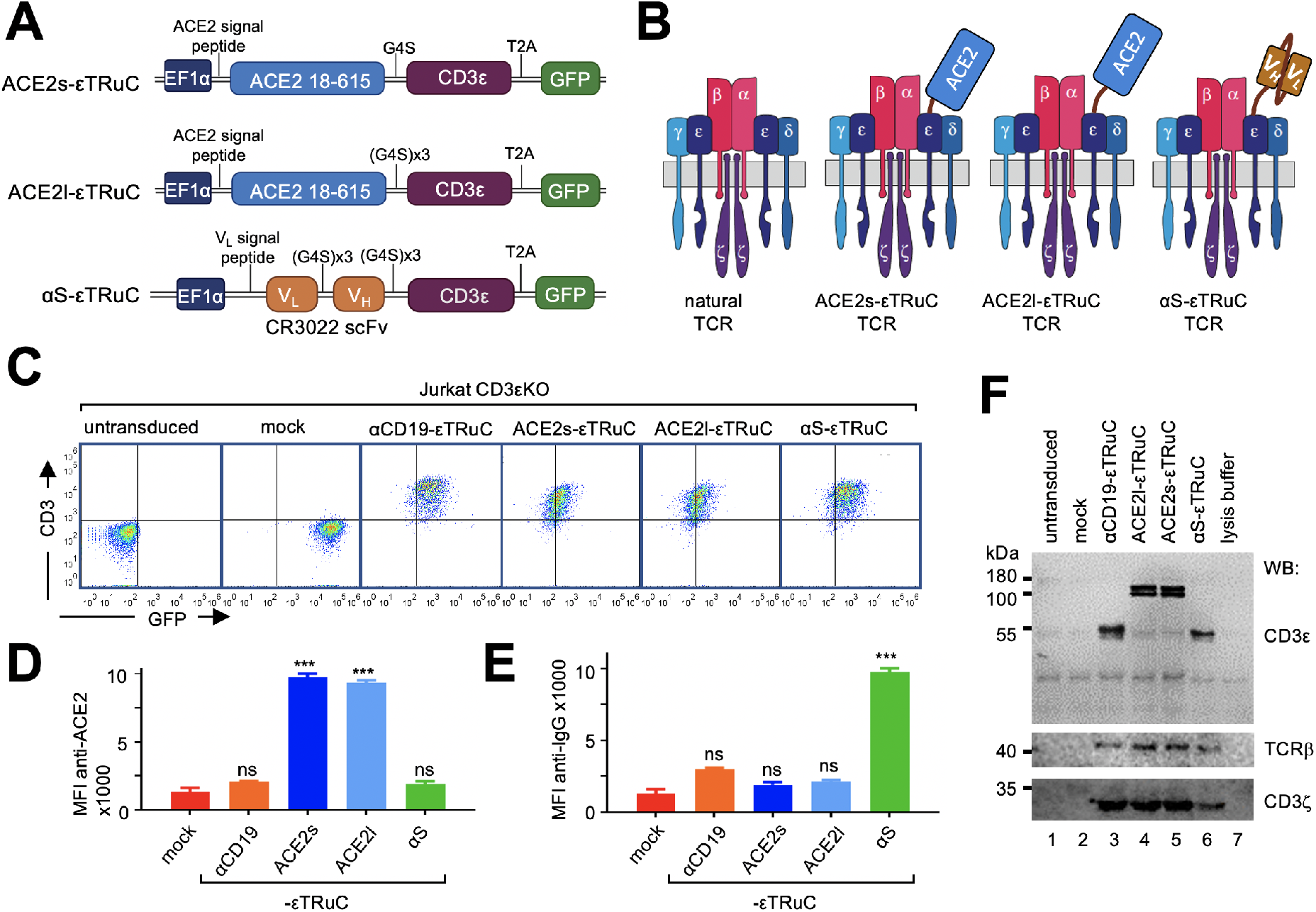
The S-specific ε TRuCs are expressed on the T cell surface as part of a TCR. **A**, Schematics of the lentiviral vectors expressing the εTRuC transgenes. **B**, The natural TCR and the three εTRuC-containing TCRs. **C**, Jurkat CD3εKO cells were transduced with the different εTRuCs or the vector encoding for GFP only (mock) or were left untransduced. Cells were stained with an anti-CD3 antibody and measured by flow cytometry (n>3). **D** and **E**, The cells from C were stained with anti-ACE2 and anti-IgG antibodies. The mean fluorescence intensity (MFI) of triplicates is shown (the experiment was repeated more than 3 times, n>3). **F**, The cells from C were lysed and TCRs immunopurified using an anti-CD3 antibody. After reducing SDS-PAGE, the CD3ε, TCRβ and CD3ζ subunits were visualized by Western blotting (n=3).

Jurkat T cells that lack CD3ε do not express a TCR on their surface (Fig. 1C, untransduced). When they were lentivirally transduced with the S-specific εTRuCs, a TCR containing these εTRuCs was expressed on the surface as seen by anti-CD3, anti-ACE2 and anti-IgG staining (Fig. 1C, D, E, S1). As controls, we used lentiviruses that only encode for GFP (mock) or for the αCD19-εTRuC. Co-purification of the endogenous TCR subunits TCRβ and CD3ζ with the εTRuCs demonstrates that the εTRuCs assemble into a TCR (Fig. 1F). Since CD3ζ is the last subunit to be added during TCR assembly (De Waal Malefyt et al., 1989; Minami et al., 1987), the data shows that the εTRuCs integrated into full TCRs. The Western blot confirmed the expected sizes of the εTRuCs (Fig. 1F). Hence, the SARS-CoV-2-specific εTRuCs are expressed as part of a TCR on the surface of Jurkat T cells.

To test whether this binding would lead to T cell activation, we used human Ramos-null B cells (with a deletion of the B cell receptor (He et al., 2018)) expressing SARS-CoV-2 S on their surface (Fig. 2A and S2A). These cells could activate our S-specific εTRuC-expressing Jurkat cells as seen by upregulation of the activation marker CD69 (Fig. 2B and S2B). Similarly, expression of the alpha (B.1.1.7) and beta (B.1.351) variants of the S protein of SARS-CoV-2, but not of the common cold human corona viruses HCoV-OC43 or −229E, led to activation of our εTRuC-expressing Jurkat cells (Fig. 2B). The S protein of SARS-CoV-1 or HCoV-NL63 weakly stimulated our T cells. This is in line with the ability of the different S proteins to bind to ACE2 and the antibody CR3022 (Delmas et al., 1992; Hofmann et al., 2005; Tian X et al., 2020; Yeager et al., 1992). As a control, Ramos cells without S did not activate these Jurkat cells. Since Ramos cells express CD19, the αCD19-εTRuC Jurkat cells were activated by all Ramos cell lines (Fig. 2B). In conclusion, Jurkat cells expressing the SARS-CoV-2-specific εTRuCs recognize and are activated by target cells expressing S from SARS-CoV-2.

**Fig. 2.**
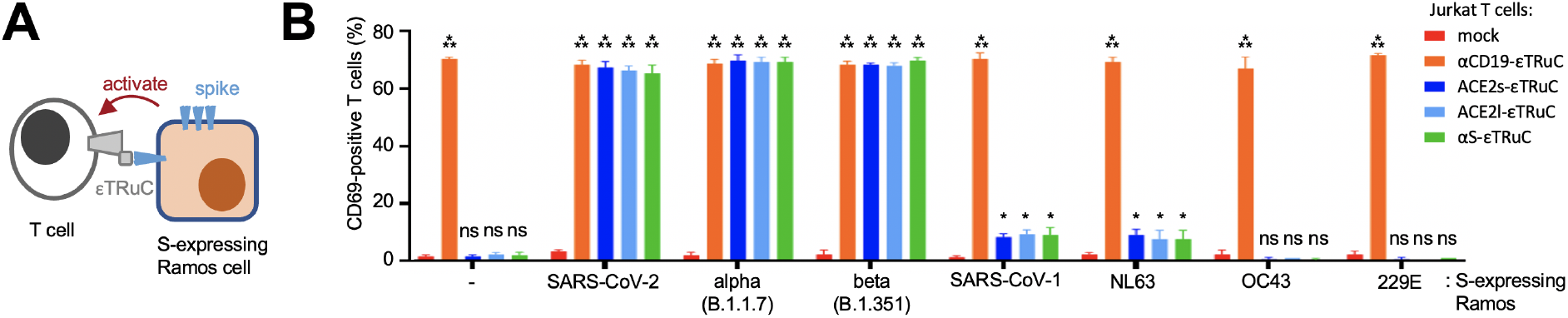
The new ε TRuCs re-program Jurkat cells to recognize S-expressing cells. **A**, By binding to the S-specific εTRuCs, Ramos cells expressing S activate the εTRuC Jurkat cells. **B**, Ramos cells without S and those expressing SARS-CoV-2 S were co-cultured with the Jurkat CD3εKO transductants from A and stained with anti-CD69 antibodies. After flow cytometric measurement, T cells were gated and the percentage of CD69-positive cells of triplicates is shown (the experiment was repeated more than 3 times, n>3).

### The novel primary ε TRuC T cells lyse S-containing target cells

Next, expanded primary human T cells were transduced to express the εTRuCs, in order to test for the cytotoxic capability of these T cells. All εTRuCs were expressed well on the surface of the T cells (Fig. 3A), which were CD8^+^ and CD4^+^ T cells (Suppl. Fig. S3A).

**Fig. 3.**
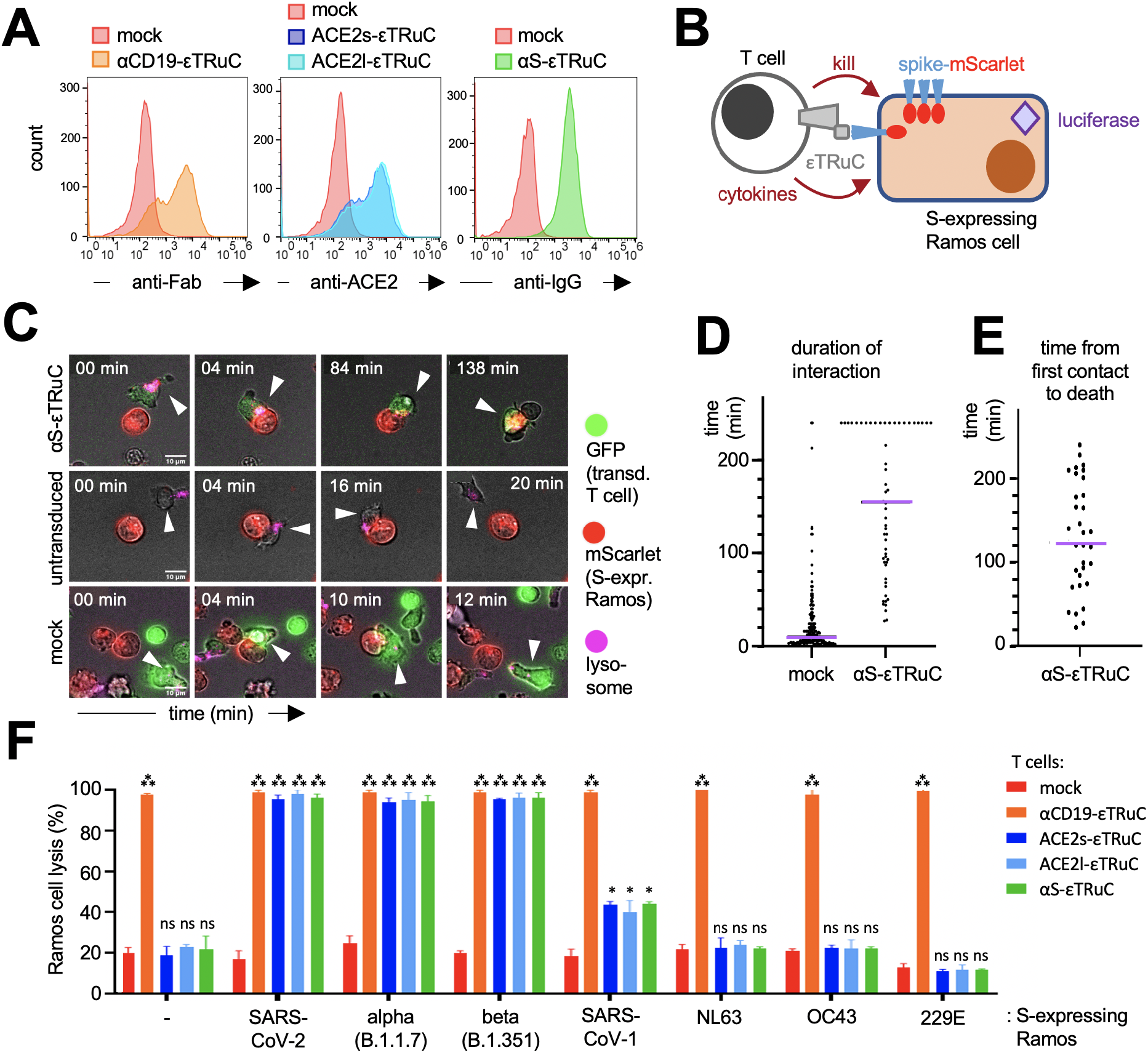
S-specific ε TRuC T cells selectively eliminate S-expressing cells. **A**, Primary human T cells were transduced with the lentiviral vectors encoding for the εTRuCs or the mock vector and expanded with IL-2. Surface expression of the εTRuCs was determined by anti-Fab, anti-ACE2 and anti-IgG staining as indicated. **B**, S-and luciferaseexpressing Ramos cells are killed by the new εTRuC T cells. **C**, An αS-εTRuC T cell (green) with the lysosomes stained in pink and an S-mScarlet-expressing Ramos cell (red) were imaged and selected frames of the given times are shown (upper panel). Non-transduced (middle panel) and mock-transduced (green, lower panel) T cells were imaged together with the S-mScarlet-expressing Ramos cells (red). **D**, Quantification of the duration of interaction between S-mScarlet-expressing Ramos cells and αS-εTRuC or mock T cells from the 4 h videos. **E**, Quantification of the time it takes for an αS-εTRuC T cell to kill an S-mScarlet-expressing Ramos cell (the purple lines in D and E depict the median). **F**, Ramos cells expressing luciferase, BFP and the different S-proteins were co-cultured with the εTRuC T cells for 24 h at in a 1:1 ratio. Target cell lysis was measured by a loss of luciferase activity in triplicates (the experiment was repeated more than 3 times, n>3).

As an example to look at target cell killing, we chose the αS-εTRuC T cells which were co-cultured with Ramos cells that express a chimeric SARS-CoV-2 S protein with a C-terminal (cytosolic) fusion to the red fluorescent protein mScarlet (S-mScarlet, Fig. 3B). Hence, transduced T cells (green) can be distinguished from the target cells (red) by confocal fluorescent imaging (Fig. 3C). After contact of the αS-εTRuC T cells with the S-expressing Ramos cells, a stable cell conjugate was formed that lasted on average 153 min (Fig. 3 D). Most likely an immunological synapse was formed as seen by the polarisation of the lysosomes (stained in magenta) towards the target cells (Fig. 3C, upper row, second image). In most cases, the S-expressing Ramos cells were killed as detected by membrane blebbing (upper row, last image) that is typical for apoptosis. When the target cell was killed, this took on average 121 min (Fig. 3E). As a control, both mock transduced T cells and untransduced T cells, had only short contact to the Ramos cells (average 10 min) and left without cytotoxic activity (Fig. 3C and D).

All S-specific εTRuC and control engineered primary T cells were co-cultured with the Ramos cells, which served as target cells, expressing or not the SARS-CoV-2 S, to quantify the cytotoxic activity. The target cells also expressed the firefly luciferase and BFP (Suppl. Fig 2C). If they were lysed, luciferase activity was lost, serving as a readout for the killing of the target cells by the T cells. Indeed, all εTRuC T cells lysed the S-expressing Ramos cells very efficiently, whereas mock T cells did not (Fig. 3F). As expected only the αCD19-εTRuC T cells killed the Ramos cells that did not express the S protein. Ramos cells expressing S of the alpha (B.1.1.7) and beta (B.1.351) variants were also lysed very efficiently, but not those expressing S from HCoV-NL63, −OC43 or −229E (Fig. 3F). There was a weak, but clear, activity also against Ramos cells expressing S from SARS-CoV-1. As above, we did not observe any differences between the ACE2l-, ACE2s- and αS-εTRuCs. Together these data demonstrate that we were able to redirect primary T cells to kill efficiently target cells that display SARS-CoV-2 S.

### The S-specific primary ε TRuC T cells are activated by S-expressing target cells

Incubation of our engineered primary T cells with the Ramos cells expressing S from different coronaviruses showed that the SARS-CoV-2 S protein led to upregulation of CD69 on the T cells (Fig. 4A). The same was observed for S of the alpha (B.1.1.7) and beta (B.1.351) variants. The ones from SARS-CoV-1 and HCoV-NL63 caused a slight CD69 upregulation, whereas the ones from HCoV-OC43 or −229E were inactive. In the T cells that are not specific for the SARS-CoV-2 S, CD69 was not upregulated by any S-expressing Ramos cell (Fig. 4A). We also show that the engineered T cells that were activated by SARS-CoV-2 S-expressing target cells secreted the cytokines IL2, IFNγ, IFNα and TNFα (Fig. 4B). As a control, the mock T cells did not produce any of those cytokines nor did a co-culture with Ramos cells that do not express any S (Fig. S4A). In conclusion, our re-programmed T cells with the new S-specific εTRuCs could be activated by S-expressing target cells.

**Fig. 4.**
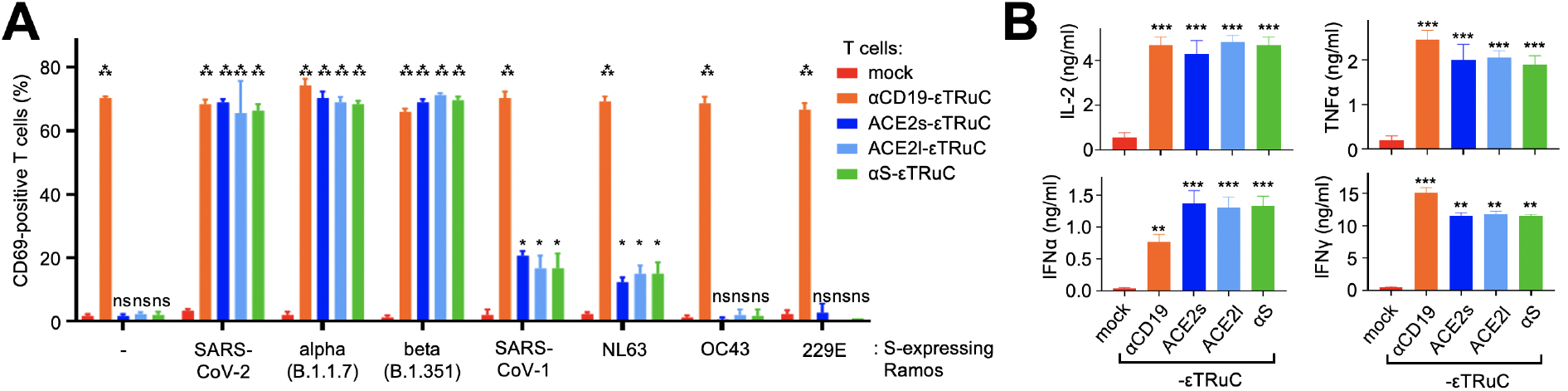
S-specific ε TRuC T cells are activated and secrete cytokines upon stimulation with S-expressing cells. **A**, Percent of CD69-positive εTRuC T cells after co-culture with the different S-expressing Ramos cells was determined by flow cytometry in triplicates. **B**, Secretion of cytokines by the εTRuC T cells following co-culture with the Ramos cells expressing SARS-CoV-2 S was quantified by ELISA in triplicates. A and B were repeated more than 3 times (n>3).

### The new S-specific primary ε TRuC T cells kill SARS-CoV-2-infected cells

Having shown that the S-specific εTRuC T cells kill S-expressing Ramos cells, we wanted to test whether these TRuC T cells are also able to kill SARS-CoV-2-infected Vero E6 cells (Fig. 5A, upper panel). To quantify killing as above, Vero E6 cells were lentivirally transduced to express luciferase and BFP bicistronically separated by an IRES sequence (called VeroLucBFP cells, Fig. S5A) and were infected with the authentic B.1 SARS-CoV-2 using a multiplicity of infection (MOI) of 0.05. After one hour, remaining virus was washed away, T cells were added and the luciferase signal was recorded (Fig. 5A, lower panel). When the T cells had been transduced with the S-specific εTRuCs (Fig. 5B, green and blue) the luciferase activity of the infected VeroLucBFP culture was reduced at 2 and 4 hours compared to the one treated with the mock and αCD19-εTRuC T cells (red and orange). The reduction of luciferase activity at early time points indicated that S-specific εTRuC T cells killed infected VeroLucBFP cells specifically, since treatment with the S-specific εTruC T cells did not cause a decrease in luciferase activity in uninfected VeroLucBFP cells beyond background (Fig. 5C). Elimination of infected cells within 2-4 hours is in line with the fast killing of Ramos cells expressing the S (Fig. 3E).

**Fig. 5.**
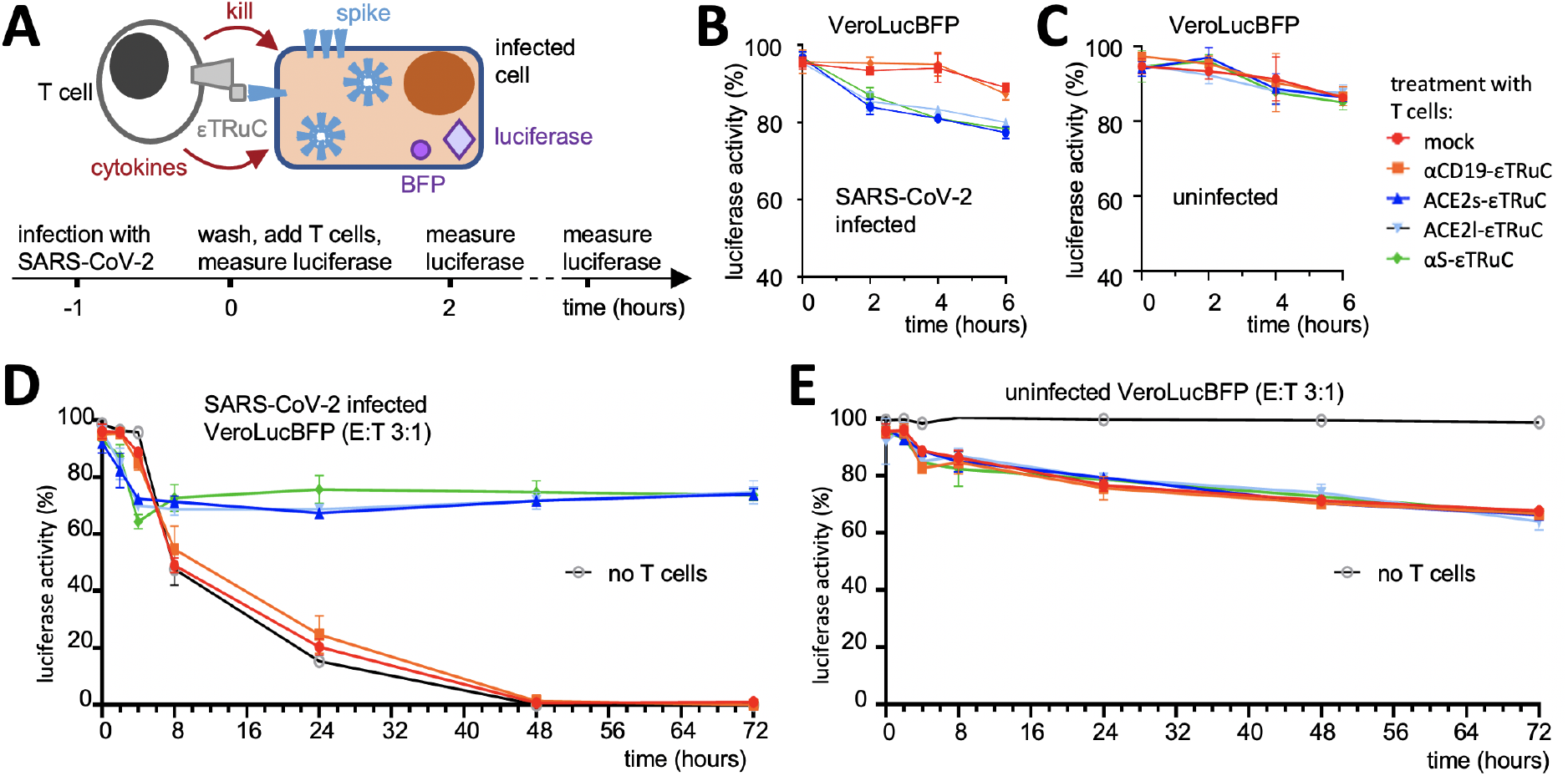
SARS-CoV-2-infecetd cells are killed by S-specific ε TRuC T cells. **A**, The new εTRuC T cells recognise and kill VeroLucBFP cells infected with SARS-CoV-2 (upper panel); scheme of the infection and T cell treatment (lower panel). **B** and **D**, VeroLucBFP cells were infected with SARS-CoV-2 at an MOI of 0.05 for 1 h. After washing the εTRuC-transduced or mock T cells were added at a T : VeroLucBFP cell ratio of 3:1 (E:T). The luciferase activity was determined for 6 h (B) or 72 h (D). Samples without the addition of T cells were included in D. Triplicates are shown. **C** and **E**, The experiments were performed as in B and D, but the VeroLucBFP cells were not infected. B to E was done with three different T cell donors. B to E were repeated more than 3 times (n>3).

Similar results were obtained when the human colorectal adenocarcinoma cell line Caco-2 or the human lung epithelial cell line Calu-3 cells were transduced with luciferase and taken as host cells for the virus infection (Fig. S5B and C). In conclusion, engineered primary human T cells expressing ACE2s-, ACE2l- and αS-εTRuCs were successfully re-programmed to recognize and kill cells infected with SARS-CoV-2.

### S-specific ε TruC T cells prevent the loss of luciferase activity in infected cells

Next, we repeated the treatment of infected VeroLucBFP cells with our T cells, but recording luciferase activity for 72 hours. With the control T cells (mock and αCD19-εTRuC) the luciferase signal started to decrease 6 hours post infection (Fig. 5D, red and orange). A similar drop in luciferase activity was observed when no T cells were added (black line), showing that T cells *per se* did not recognize SARS-CoV-2-infected cells. At 72 hours post infection no luciferase activity was detected anymore. This loss of luciferase signal was prevented by the presence of the S-specific εTRuC T cells (green and blue). In fact, at 48 and 72 hours post infection the infected and treated cultures had similar luciferase activity as the uninfected ones (Fig. 5D and E), demonstrating that the TRuC treatment resulted in a sustained luciferase expression.

Similar results were obtained when a different ratio of T cell to VeroLucBFP cells was used (Fig. S6) or when Caco-2 or Calu-3 cells were infected (Fig. S6). In all cases, our treatment completely rescued the infected cell cultures in terms of the virus-induced loss of luciferase activity.

### SARS-CoV-2 causes a translation shutdown in VeroLucBFP cells

In theory, the loss of luciferase signal may result from three different mechanisms. (i) By killing of VeroLucBFP cells by the S-specific TRuC T cells (as above). (ii) By reducing the VeroLucBFP cell number by either SARS-CoV-2-induced cell death (Park et al., 2020) or reduced cellular proliferation rates (our graphs depict the luciferase activity of the wells in relation to the wells with the uninfected cells). (iii) By the SARS-CoV-2-mediated suppression of host protein synthesis (Hsu et al., 2021; Lapointe et al., 2021; Puray-Chavez et al., 2020; Thoms et al., 2020; Yuan et al., 2020b), called translation shutdown. This mechanism acts at the single cell level.

To monitor the effect of translation shutdown, we blocked protein synthesis in VeroLucBFP cells with puromycin, which causes premature chain termination during mRNA translation (Pestka, 1971). Indeed, puromycin-treatment suppressed luciferase activity before the occurrence of cell death (Fig. 6A). To directly examine whether a strong translation shutdown in infected cells occurred, we determined the BFP expression in SARS-CoV-2-infeceted VeroLucBFP cells. BFP allows us to investigate the translation shutdown at the single cell level by flow cytometry. Indeed, at 72 hours post infection when no luciferase activity was detectable (Fig 5D), the BFP expression was similarly lost (in the control cultures that were treated with mock or αCD19-εTRuCs T cells) (Fig. 6B). This suggests that a complete translation shutdown is induced by SARS-CoV-2 in the VeroLucBFP cells, as seen earlier in other cells (Hsu et al., 2021; Lapointe et al., 2021; Puray-Chavez et al., 2020; Thoms et al., 2020; Yuan et al., 2020b). This loss of BFP-expression was not observed in uninfected VeroLucBFP cells (Fig. 6B and S5A). SARS-CoV-2 infection also led to slightly reduced VeroLucBFP cell numbers at 72 hours (Fig. S7A), most likely due to viral lysis of the host cells (Park et al., 2020).

**Fig. 6.**
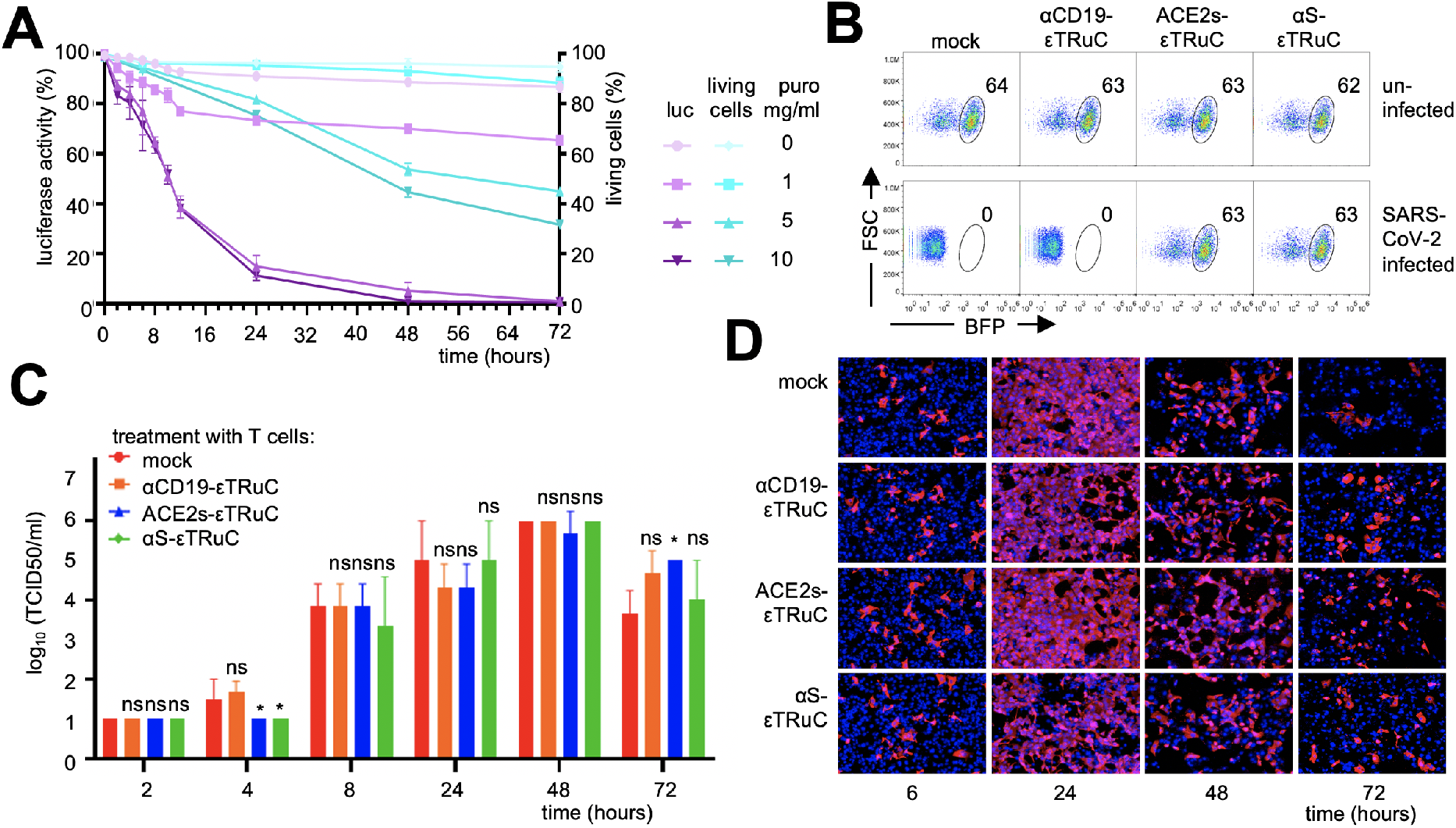
ε TRuC T cell treatment prevents the protein shutdown in VeroLucBFP cells, but does not limit viral replication. **A**, VeroLucBFP cells were treated with puromycin (0-10 μg/ml). The luciferase activity (purple lines) and the percent of living cells (turquoise lines) were determined. **B**, VeroLucBFP cells were infected with SARS-CoV-2 (MOI 0.05) for 1 h or left uninfected. After washing the transduced T cells were added at a ratio of T : VeroLucBFP cell of 3:1 and at 72 h post infection the BFP-expression of the VeroLucBFP cells was quantified by flow cytometry. The percent of BFP^+^ cells is given. **C** and **D**, In an experiment as B, the viral titers in the supernatant of the infected and treated VeroLucBFP cells (C) and the amount of infected cells as seen by staining for the viral N protein in pink (D) was determined at several time points post infection. In D, the nuclei of the cells were stained with DAPI (blue).

### S-specific ε TRuC T cells prevent the translation shutdown, but not viral replication

The loss of luciferase activity at later time points post infection, which is indicative for the translation shutdown, was drastically reduced upon S-specific εTRuC T cell treatment (Fig. 5D and S6). Indeed, treatment of infected VeroLucBFP cells with ACE2s- or αS-εTRuC T cells, also prevented the loss of BFP expression (Fig. 6B). The infected and treated VeroLucBFP cells had similar BFP values as the uninfected ones. Assuming that the S-specific εTRuC-treatment would prevent the translation shutdown by an killing of all infected VeroLucBFP cells immediately post infection, we expected no viral replication and spread in the tissue culture. Surprisingly, although some killing of infected cells was detected at early time points (Fig. 5B and S5), SARS-CoV-2 replicated equally well independent of our treatment. This was seen by determining the virus titer of the culture supernatants at different time points post infection (Fig. 6C) and by SARS-CoV-2 nucleocapsid protein (N) staining in infected cells (Fig. 6E). In conclusion, at later time points post infection (8 to 72 hours) the virus replicates, infects new VeroLucBFP cells and leads to a suppression of the host protein synthesis in the VeroLucBFP cell culture. Importantly, this shutdown was completely prevented by treatment with our engineered T cells.

Lastly, we sought to get insight into how the S-specific εTRuC T cell blocks the virus-induced translation shutdown. To this end, we activated ACE2s- or αS-εTRuC T cells by co-culturing them with S-expressing Ramos cells for 24 hours. We then collected the medium that should contain factors that these T cells secrete and treated uninfected and SARS-CoV-2-infected VeroLucBFP cells with this “conditioned” medium. Upon treatment of SARS-CoV-2-infected cells with the ACE2s- or αS-εTRuC T cell “conditioned” medium, the loss of luciferase activity and of BFP expression was reduced compared to cells incubated with “conditioned” medium derived from mock T cells (Fig. 7A and 7B). These data suggest that our engineered T cells prevent virus-induced protein shutdown in VeroLucBFP cells by secreting a protective factor or a cocktail of different factors instead of killing infected cells. This is in line with the observation that the T cells also prevented the translation shutdown, when using a low MOI, where the killing of infected cells at the early time point was not visible (Fig. S7B).

**Fig. 7.**
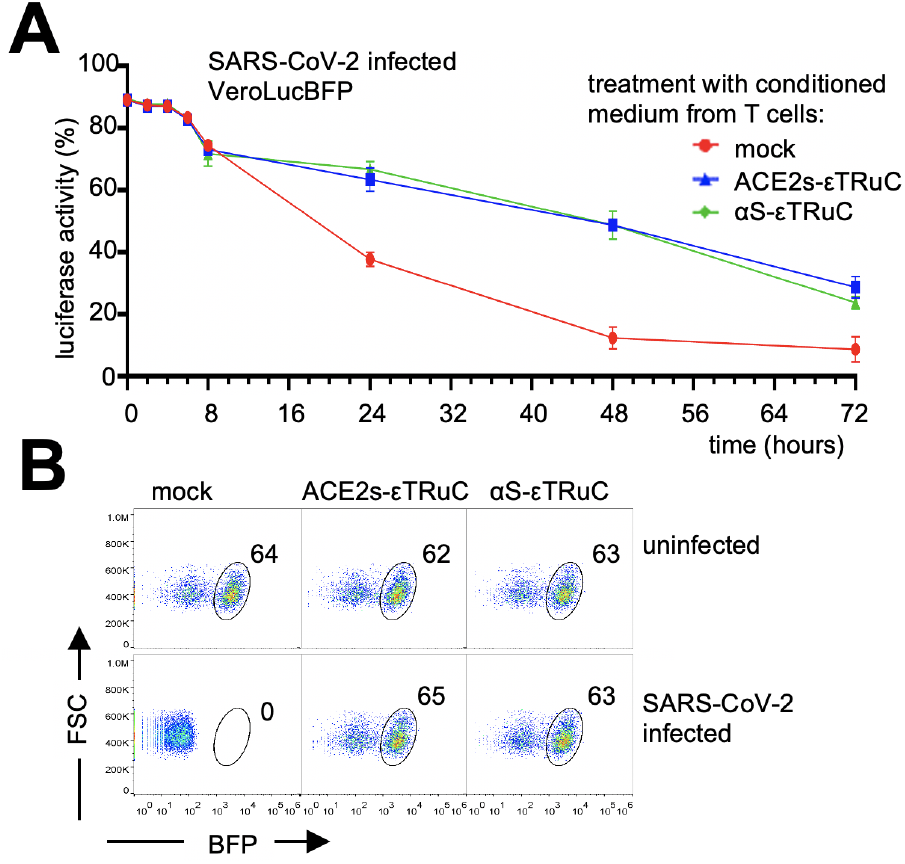
Soluble factors of the εTRuC T cells prevent the protein shutdown in VeroLucBFP cells. **A**, Conditioned medium of mock, ACE2s- and αS-εTRuC T cells that were co-cultured with S-expressing Ramos cells was added to SARS-CoV-2-infected VeroLucBFP cells at 0 and again at 24 h post infection. Luciferase activity was measured in triplicates (the experiment was done twice, n=2). **B**, At 72 h post infection the BFP-expression of the VeroLucBFP cells treated with the conditioned medium was quantified by flow cytometry. The percent of BFP^+^ cells is given.

## Discussion

Our results demonstrate that the TRuC technology can be used to re-program T cells to recognize SARS-CoV-2-infected cells in an HLA-independent manner. TRuC T cells have been developed against tumor cells, now widening their potential to viral infections, such as Covid-19. Instead of recognizing a tumor antigen, the new εTRuCs recognize the SARS-CoV-2 S protein, and thus distinguish healthy from infected cells. In fact, the here presented εTRuCs were generated by either fusing the S-binding part of the ectodomain of ACE2 (Lan et al., 2020; Yan et al., 2020; Yuan et al., 2020a; Zhou et al., 2020) or the scFv of the anti-S antibody CR3022 (ter Meulen et al., 2006; Wrapp et al., 2020; Yuan et al., 2020a) to the TCR,s CD3ε chain. The resulting S-specific εTRuCs integrated into complete TCRs, as seen by surface expression of the εTRuC (in wild-type and CD3ε KO T cells) and co-immunoprecipitation of endogenous TCR subunits. Importantly, the engineered T cells specifically lysed SARS-CoV-2-infected cells and secreted anti-viral cytokines.

Ramos tumor cells exogenously expressing SARS-CoV-2 S on their surface were lysed as efficiently by our novel εTRuC T cells as by the αCD19-εTRuC T cells targeting the tumor antigen CD19 that is also expressed by Ramos. Since the αCD19-εTRuC T cells are very potent against B cell-derived tumors (Baeuerle et al., 2019), the new S-specific εTRuCs demonstrated full functionality. The observed killing events were very fast (between 30 min and 4 hours) and included recognition of the target cells as seen by prolonged conjugate formation. Ramos cells expressing S from the alpha (B.1.1.7) and beta (B.1.351) variants were also killed efficiently, suggesting that these currently circulating variants of concern might be equally well targeted by our εTRuCs. Importantly, while escape from neutralising antibody binding is frequently observed for SARS-CoV-2 variants (Hoffmann et al., 2021), the reliance on ACE2-mediated entry into the host cells by SARS-CoV-2 suggests that escape from binding to ACE2-εTRuC will impose an unlikely strong fitness cost. Ramos cells expressing S from SARS-CoV-1 and HCoV-NL63 were recognized to a certain degree, since S of these viruses also binds to ACE2, albeit with lower affinity (Horndler et al., 2021; Wu et al., 2009; Xie et al., 2020). Ramos expressing S from HCoV-OC43 or −229E that do not bind to ACE2 (Delmas et al., 1992; Yeager et al., 1992) were not killed, showing again the specificity of our approach. Together our data are in line with two recent unreviewed preprints. They report that expressing S-specific CARs based on the antibodies CR3022 or S309 in natural killer (NK) cells led to the killing of S-expressing target cells (Ma et al., 2021; Ma et al., 2020).

Cytotoxic activity of our εTRuC T cells was also seen against SARS-CoV-2-infected Vero E6, CaCo-2 and Calu-3 cells, indicating that the amount of S being displayed on the surface of the infected cells unleashed recognition and killing by the novel εTRuC T cells. Indeed, T cell activation via the TCR can be mediated by very low amounts of antigen, such as 3-7 copies per target cell (Irvine et al., 2002; Purbhoo et al., 2004). Lysis of the infected cells occurred within 2 hours post infection, suggesting that the post-fusion and membrane remaining S protein, rather than newly produced S by the host cell was recognized. However, the latter possibility cannot be ruled out and indeed the wild-type S protein can reach the cell surface when exogenously expressed (Horndler et al., 2021; Lapuente et al., 2021).

Although infected cells were killed quicker than the complete virus replication cycle takes (which is around 8 hours), the virus could replicate and spread within the cultures. Hence, either not all infected cells were killed or killing occurred too late, so that virions were already formed. This is a clear difference in the usage of εTRuC T cells targeting tumor cells, since in the latter case, it does not matter if killing takes longer.

SARS-CoV-2 induces a stop in translation of endogenous proteins by the infected cells (Banerjee et al., 2020; Hsu et al., 2021; Lapointe et al., 2021; Puray-Chavez et al., 2020; Thoms et al., 2020; Yuan et al., 2020b). In fact, it was shown that the viral protein Nsp1 blocks the ribosome (Thoms et al., 2020; Yuan et al., 2020b) and in our experiments this was detected by both a loss of luciferase activity and a loss of BFP expression at the individual cell level. Puromycin, which also prevents protein translation, led to a loss of luciferase activity too and this occurred earlier than cell death, indicating again that luciferase activity is a read out for the translation shutdown in our experiments. Indeed, luciferase activity and fluorescence of a reporter (GFP) were used earlier to quantify this shutdown (Hsu et al., 2021; Lei et al., 2020; Puray-Chavez et al., 2020; Thoms et al., 2020; Xia H et al., 2020; Yuen et al., 2020).

Treatment of the infected cultures with the S-specific εTRuC T cells completely prevented the virus-induced translation shutdown; and 72 hours post infection infected and treated cultures resembled the non-infected controls in terms of luciferase activity and BFP expression. Thus, the engineered T cells provided long-term protection from the translational arrest. This effect of the T cells was robustly seen with various MOIs and T cell:infected cell ratios. A reduction in the shutdown was also evident when only the supernatant of activated εTRuC T cell was used, suggesting that one or more soluble factor(s) mediated this protective effect. Thus, our hypothesis is that upon recognition of infected cells, the S-specific εTRuC T cells are activated (as seen by the killing in the first hours) and secrete cytokines or other factors that can rescue the infected cells from the protein synthesis shutdown. Indeed, we show that these T cells produce certain cytokines, such as INFγ, IFNα, TNFα and IL-2, upon stimulation with S-expressing Ramos cells. However, the identity of the factor counteracting the shutdown upon infection remains enigmatic.

With or without translation shutdown the same amount of virus was detected in the supernatant of the infected Vero E6 cells, indicating that a stop in the translation of host proteins does not impair viral growth. Thus, the main aim of the shutdown might be to prevent the innate anti-viral response by the host cell, as suggested earlier (Lei et al., 2020; Puray-Chavez et al., 2020; Thoms et al., 2020; Xia H et al., 2020; Yuen et al., 2020). Importantly, by preventing this shutdown our treatment might enable a potent anti-viral response, helping in the noncytolytic clearance of the virus, thereby preventing strong tissue damage. The latter is relevant, because in some viral infections (such as in the lymphocytic choriomeningitis or hepatitis B viruses (Guidotti et al., 1999; Guidotti and Chisari, 2006)) most of the cells in a certain organ are infected and massive killing of the infected cells might destroy the organ. Surprisingly, although the S-specific εTRuC T cells killed infected cells in the first hours, they spared infected cells later on and allowed those to survive and prevent the translation shutdown.

CARs have been successfully used against hematological but not solid tumors. In contrast, εTRuC T cells are active against solid tumors and show enhanced efficacy against hematological malignancies (Baeuerle et al., 2019). This is most likely due to the employment of the full TCR signal, in contrast to the CARs that only use part of the TCR signal (namely the one by the isolated CD3ζ chain). Furthermore, the strength of the TCR (and TRuC-TCR) signal is autoregulated and finely balanced by conformational changes of the full TCR complex (Gil et al., 2002; Hartl et al., 2020; Minguet et al., 2007; Swamy et al., 2016). For example, the strong tonic signaling by the CARs that might lead to CAR T cell exhaustion (Ajina and Maher, 2018; Frigault et al., 2015) is not seen with the εTRuCs. Lastly, εTRuC expression is limited by its incorporation into the TCR, whereas CARs are often overexpressed (Baeuerle et al., 2019). Thus, CAR T cells cannot benefit from 400 million years of TCR evolution.

In conclusion, with our εTRuC technology, derived from the cancer field, we re-programmed human T cells to specifically recognize SARS-CoV-2-infected cells. This might be the basis to develop an effective treatment for severely diseased Covid-19 patients.

## Methods

### Molecular cloning of the ε TRuC constructs

The plasmids used in this study were generated using standard molecular cloning techniques, such as PCR, restriction digest and Gibson assembly (Gibson et al., 2009). pOSY120, encoding for the ACE2l-εTRuC, was generated by Gibson assembly of the XhoI-XbaI fragment of p526 anti-αCD19 (Baeuerle et al., 2019) and the PCR fragment using the primers O293 and O294 (see table 1) on the human ACE2 sequence as a template. pOSY121, encoding for the ACE2s-εTRuC, was generated by Gibson assembly of the XhoI-XbaI fragment of above and the PCR fragment using the primers O293 and O295 on the human ACE2 sequence. pOSY123, encoding the αS-εTRuC, was generated by Gibson assembly of the XhoI-XbaI fragment of above and, the PCR fragment using the primers O304 and O305 from a gBlock of CR3022 V_L_ (Integrated DNA Technologies) and the PCR fragment using O306 and O307 from a gBlock of CR3022 V_H_ (Integrated DNA Technologies). All plasmid sequences were verified by Sanger sequencing (Eurofins Genomics).

**Table 1.**
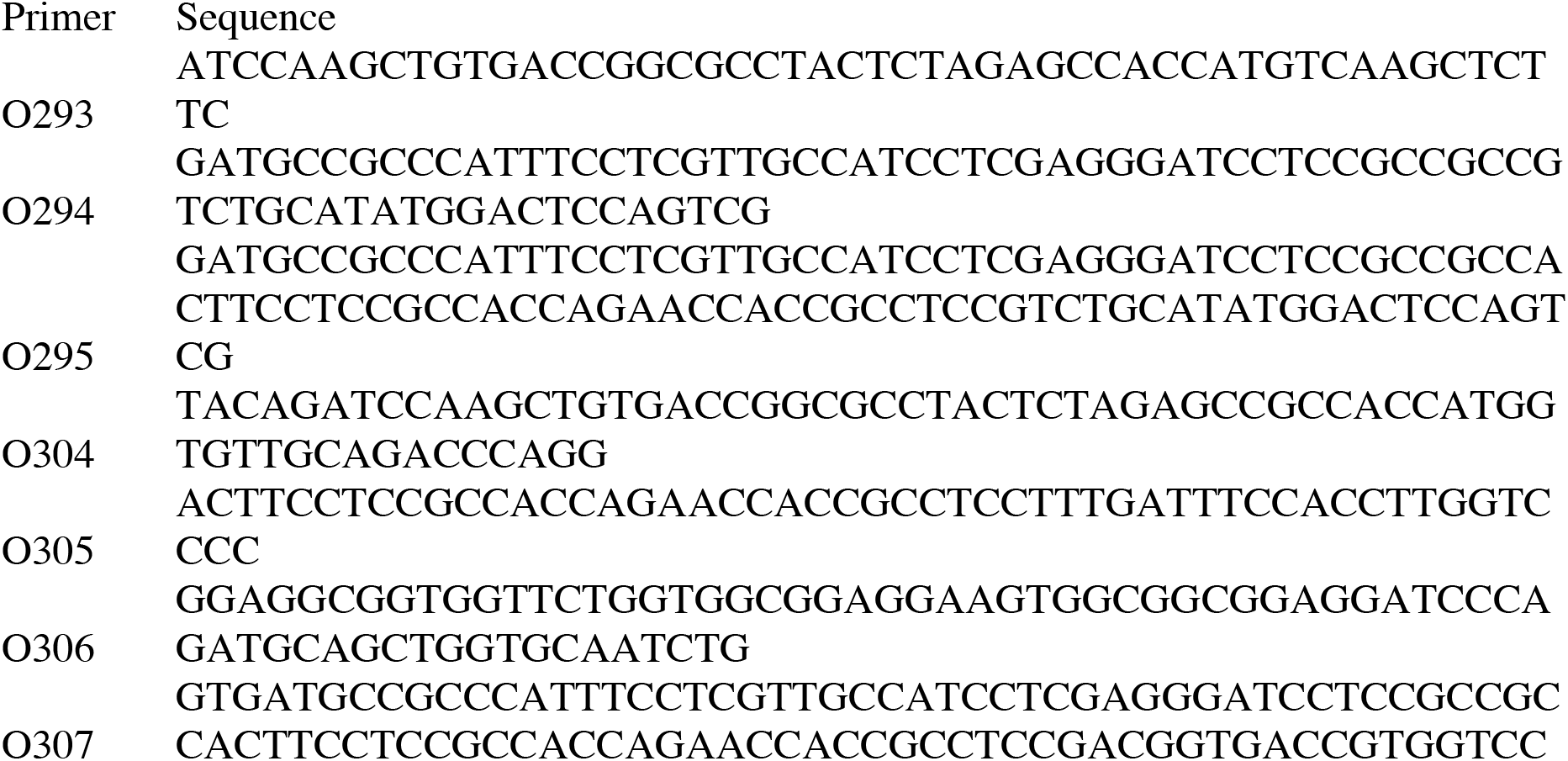
Primer sequences

### Molecular cloning of the luciferase-BFP and S protein-mScarlet vectors

For the molecular cloning of the pHRSIN-CS-Luc-IRES-mTagBFP2 vector, the previously described pHRSIN-CS-Luc-IRES-emGFP vector (a kind gift from A. Rodriguez, Universidad Autonoma Madrid, Spain) was digested with BstX1 and Not1 restriction enzymes to remove the emGFP. mTagBFP2 was amplified from the previously described pHRSIN-CS-IRES-mTagBFP2 plasmid (Dang et al., 2020) and BstX1- and Not1-specific overhangs were added using PCR. The final expression vector was generated by Gibson assembly using the pHRSIN-CS-Luc-IRES as recipient vector and the amplified mTagBFP2 as insert. The integrity of the plasmid was verified by Sanger sequencing.

The retroviral expression vector encoding the S protein of SARS-CoV-2 fused at is cytoplasmic tail to mScarlet was cloned as follows. The cDNA of the S protein was taken from the plasmid pCG1-CoV-2019-S with a codon-optimized sequence (Lapuente et al., 2021) and cloned into the pMIG vector by Gibson assembly and the cDNA of mScarlet was ligated into the vector.

All plasmid sequences were verified by Sanger sequencing (Eurofins Genomics).

### Generation of lentiviruses

Lentiviruses were prepared as described (Hartl et al., 2020). In brief, HEK293T cells were transfected with the εTRuC-encoding lentiviral plasmids and the packaging plasmids pMD2.G (envelope) and pCMVR8.74 (gag/pol) using PEI transfection. The virus-containing supernatant was collected, concentrated by centrifugation and the virus pellet was resuspended in RPMI 1640 medium and stored at −80°C. If MOI is explained in Frederikes Nat Immunol paper, then we can cite this here. If not we need to write a few sentences.

### Cells

Jurkat (human T cell line), Ramos (human Burkitt’s lymphoma line), VeroE6 (kidney epithelial cell line form the African green monkey), CaCo-2 (human colorectal adenocarcinoma line) and CaLu-3 (human lung epithelial cell line) cells and their derivatives were grown with complete RPMI 1640 medium and 10% fetal calf serum (FCS) at 37°C and 5% CO_2_. Jurkat CD3ε knock out (KO) cells were generated by standard CRISR/Cas9 technology and the generation of the Ramos cells expressing the different S proteins will be described elsewhere. Jukat CD3ε KO, Ramos, Vero E6, CaCo-2 and CaLu-3 cell lines were transduced with a multiplicity of infection (MOI) of 5 with the lentiviruses indicted and sorted by flow cytometry when necessary.

To obtain the expanded human T cells, peripheral blood mononuclear cells (PBMCs) were isolated from blood of a healthy donors by density-gradient centrifugation and grown in RPMI 1640 medium supplemented with 10% FCS and 1000 U/ml recombinant IL-2 (PeproTech) and activated with 1 μg/ml anti-CD3 and anti-CD28 antibodies. At 48-72 h the remaining PBMCs were mostly T cells (> 99%) and lentivirally transduced by spin infection with 5 μg/ml of protamine sulfate with a MOI of 4. Transduced T cells were tested using flow cytometry by staining with anti-Fab for the αCD19-εTRuC, anti-ACE2 for the ACE2s- and ACE2l-εTRuCs, and anti-human IgG for the αS-εTRuC. Cells were expanded in complete RPMI 1640 medium with 10% FCS and were given 100 U/ml IL-2 every third day. Cells were used until day 17 post transduction.

To generate S-mScarlet expressing Ramos cells, the retroviral vector pMIG encoding S-mScarlet and a vector encoding the ecotropic packaging protein were co-transfected (500 ng each) into Plat-E cells with the PolyJet transfection reagent (Signagen). After two days, the supernatant was collected, filtered and mixed 1:1 with 300.000 Ramos-null cells that express the ecotropic receptor. Cells were sorted for mScarlet expression.

The transduction of Ramos cells with LucBFP was done as follows. Retroviral transduction of the human Ramos-null B cells (with a deletion of the B cell receptor (He et al., 2018)) with vectors encoding for the different S proteins will be published elsewhere. These Ramos-null cells expressing S proteins or not were lentivirally transduced with pHRSIN-CS-Luc-IRES-mTagBFP2 as briefly described above.

### Flow cytometry

The following antibodies were used for flow cytometry staining in a 96-well format: PE-labelled anti-human CD4 (Beckman Coulter, #A07752), APC-labelled anti-human CD8 (Beckman Coulter, #IM2469), APC- or PacificBlue-labelled anti-human CD3 (BioLegend, #300434), PE-labelled anti-human CD69 (Life Technologies, #MHCD6904), anti-human ACE2 (R&D Systems, #HK0320042), PE-labelled anti-human IgG (Southern Biotech, #2040-09) and biotin-labelled anti-Fab (Invitrogen, #31803). APC-coupled Streptavidin (Biolegend, #105213) and APC-labelled donkey anti-goat IgG (Southern Biotech, #6420-05) served as a secondary reagent. Cells were measured on the flow cytometer Attune NxT and the data were analyzed by FlowJo.

For flow cytometry of virus-infected cells post-treatment, the cells were fixed with methanol and paraformaldehyde and then washed 4 times before measuring on AttuneNxT.

### Co-immunopurification and Western blotting

Following antibodies were used for biochemical analysis: anti-TCRα (clone H-1, Santa Cruz, #sc-515719), anti-TCRβ (clone H-197, Santa Cruz, #sc-9101), anti-CD3γ (clone EPR4517, Epitomics, #3256-1), anti-CD3δ (clone F-1, Santa Cruz, #sc-137137,), anti-CD3ε (clone M20, Santa Cruz, #sc-1127), anti-CD3ζ (serum 449), horseradish peroxidase (HRPO)-coupled anti-mouse IgG (Thermo Fisher, #32430), HRPO-coupled anti-goat IgG (Thermo Fisher, #31402), and HRPO-coupled anti-rabbit IgG (Thermo Fisher, #31460). Protein G-coupled sepharose (#17–0618–01) and Protein A-coupled sepharose (#17–5138–01) beads were from GE Healthcare and the protease inhibitor cocktail was from Sigma.

3 × 10^7^ cells were lysed in 0.4 ml lysis buffer containing 20 mM Tris-HCl pH8, 137 mM NaCl, 2 mM EDTA, 10% glycerol, 1x protease inhibitor cocktail, 1 mM PMSF, 5 mM iodoacetamide, 0.5 mM sodium orthovanadate, 1 mM NaF, and 0.5% Brij96 for 30 min at 4 °C followed by 15 min centrifugation to pellet the nuclei and insoluble material. For the anti-CD3ε immunoprecipitation 370 μl cleared cell lysate was incubated with 5 μl 50% protein A and protein G sepharose slurry (1:1) and 2 μg anti-CD3ε UCHT1 for 4 h at 4 °C. After three washes, the immunoprecipitated material was separated by 12% reducing SDS-PAGE. The separated proteins were transferred to PVDF membranes by semi-dry transfer. After blocking with 5% milk in PBS containing 0.1% Tween-20 the membranes were incubated with antibodies against TCRα (1:1000), TCRβ (1:100), CD3γ (1:1000), CD3δ (1:100), CD3ε (1:1000), CD3ζ (1:1000) in PBS-T followed by incubation with HRPO-conjugated secondary antibodies (1:10000). Western blot signals were recorded using an Image Quant LAS 4000 Mini from GE Healthcare Life Sciences, Boston, MA.

### Activation assays

Ramos cells expressing or not the different S proteins were co-cultured with the different Jurkat transductant cells at a 1:3 target-to-effector ratio for 9 h. Cells were stained with anti-CD69 antibodies and measured by flow cytometry. The BFP-positive Ramos cells were gated out to ensure that only the T cells are analysed.

To quantify cytokines by ELISA, εTRuC-expressing primary T cells and S-expressing Ramos cells were co-cultured for 24 h. The following cytokines were measured according to the instrzctions of each ELISA kit: TNFα (Invitrogen, #88-7346-88), IFNγ (Invitrogen, #88-7316-88), IFNα (Invitrogen, #BMS216) and IL2 (Invitrogen, #88-7025-88).

### Cytotoxicity assay

To test for the T cell-mediated lysis of the Ramos target cells expressing firefly luciferase and the S proteins indicated, a bioluminescence-based cytotoxicity assay was performed. 100.000 Ramos cells were plated in 100 μl of complete RPMI 1640 medium with 10% FCS in a 96-well flat bottom plate (Corning). 75 μg/ml D-firefly luciferin potassium salt (Biosynth) was added and bioluminescence was measured in the luminometer (Tecan infinity M200 Pro) to establish a baseline. Then, 300.000 εTRuC-expressing T cells (effector cells) were added at an effector-to-target ratio of 3:1 and incubated for 24 h at 37°C. Relative light units (RLUs) signals from target cells treated with 1% Triton X-100 indicate maximal cell death. RLU signals from target cells without added T cells determine spontaneous cell death. The percent of specific lysis was calculated with the following formula:

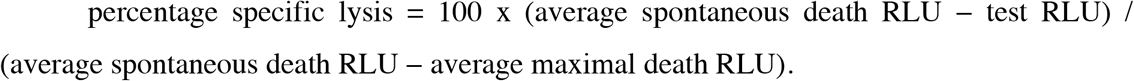

### Time-lapse microscopy

Ramos B cells were retrovirally transduced with a construct encoding spike protein tagged with the fluorophore mScarlet at the C-terminus and sorted by FACS (Bio-Rad S3e Cell Sorter). For the experiment, mScarlet-spike-expressing Ramos B cells were washed 3 times in PBS. 15,000 cells in PBS were seeded into both wells of a 2 well-culture insert placed on a 35 mm dish (Ibidi). PBS facilitated the attachment of Ramos B cells onto the ibi-treated dish surface. T cells were incubated with 50 nM Lysotracker Deep Red (Invitrogen) for 15 min at a 37°C incubator with 5% CO_2_, washed twice and resuspended in phenol red-free RPMI containing 2 mM GlutaMAX (Gibco), 10% FCS (PAN), 50 U/ml Pen-Strep (Gibco) and 10 mM HEPES (Gibco). 7,500 mock- and anti-spike-transduced T cells were seeded into the 2 well-culture insert with Ramos B cells after removing the PBS. Mock- and anti-spike-transduced T and Ramos B cell interactions were recorded in parallel using a Zeiss Observer microscope equipped with a 40x oil objective and Zen Blue software. Single plane images in 4 channels (GFP, mScarlet, Lysotracker deep red, brightfield) were acquired every 2 or 3 min for 4 h within an incubator set to 37°C and 5% CO_2_. Multi-position videos were converted to TIFF and transferred to ImageJ. Duration of interaction between T and B cells and the time it takes for an anti-spike-transduced T cell to kill mScarlet-spike-expressing Ramos B cell were quantified manually. Cell death was assessed by apoptotic morphology and the subsequent halt in cellular movement from the brightfield images.

### SARS-CoV-2 infection and measurement of luciferase activity

A bioluminescence based cytotoxicity assay was performed with transduced primary cells co-cultured with VeroE6 (kidney epithelial cells extracted from an African green monkey), CaCo-2 (human colorectal adenocarcinoma cells) and CaLu-3 (human lung cancer cell line) cell lines. Each of these cell lines were infected with the B.1 SARS-CoV-2 virus at the indicated MOI. A control setup had the same cell lines without the infection. 10^4^/μl of target (Vero E6, CaCo-2 and CaLu-3) cells were plated in a 96-well flat bottom plate (Corning, #3917). 75 μg/ml D-firefly luciferin potassium salt (Biosynth) was added to it and bioluminescence (BLI) was measured in the luminometer (Tecan infinity M200 Pro) to establish the BLI baseline. Right after, TRuC-expressing T cells (effector cells) were added at varying effector-to-target ratio (as indicated) and incubated until 72 h post infection (as indicated) at 37°C. BLI was measured as relative light units (RLUs). RLU signals from cells treated with 1% Triton X-100 indicate maximal cell death. RLU signals from tumor cells without TRuC T cells determine spontaneous cell death. Percent specific lysis (specific cytotoxicity) was calculated with the following formula:

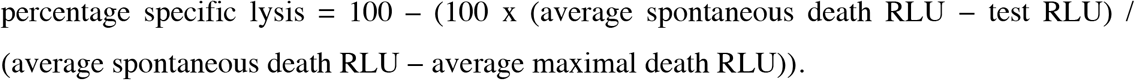

Here, spontaneous death was considered of only the target cells without the virus and without T-cells.

### Immunofluorescence and viral titers

Virus containing supernatant was harvested at the indicated time points and the viral titer was determined on VeroLucBFP cells by indirect-immunofluorescence. Briefly, VeroLucBFP cells were seeded in 96-well plates at a density of 0.04 x 10^6^ cells/well 24 h prior to infection. Infectious cell supernatants were diluted in tenfold dilution series in PBS containing 2% BSA in a 100 μl volume and subsequently incubated on VeroLucBFP. At 20 h post infection, the infectious supernatant was removed and cells were fixed using 4% formaldehyde for 30 minutes. Virus-infected cells were subsequently detected using anti-SARS-CoV nucleocapsid (N) rabbit antiserum (Rockland Immunochemicals, #200-401-A50) and secondary anti-rabbit IgG-coupled to Cyanine Cy3 (Jackson ImmunoResearch). Nuclei were stained with DAPI. Viral endpoint titers were evaluated by fluorescence microscopy.

To monitor SARS-CoV-2 spread upon εTRuC T cell-treatment, infected VeroLucBFP cells were fixed at the indicated time points post infection with 4% formaldehyde for 30 minutes and subjected to indirect-immunofluorescence as described. Fluorescence images were acquired using a Zeiss Observer.Z1 inverted epifluorescence microscope (Carl Zeiss) equipped with an AxioCamMR3 camera using a ×20 objective.

### Statistics

Statistical significance of one sample in comparison to another (a control) was determined using a paired Student,s t-test. All p values indicated with stars (* < 0.05, ** < 0.005, *** < 0. 0005) were calculated using Prism v.6 software (GraphPad). Error bars show standard deviation in all graphs.

## Acknowledgements

We thank Stephanie Pfändeμ Bochum, Germany, and Susana Minguet, Freiburg, Germany, for their expert input, and TCR^2^ Therapeutics, Cambridge, USA, for the αCD19-εTRuC plasmid and Matthias Tenbusch, Erlangen, Germany, for the S protein plasmid. We thank Simone Lölhöffel von Löwensprung for technical help. This study was supported by the German Research Foundation (DFG) through BIOSS - EXC294 and CIBSS - EXC 2189 to W.W.S., SFB854 (B19 to W.W.S.), SFB1381 (A9 to W.W.S.) as well as FOR2799 (SCHA 976/8-1) and SCHA 976/7-1, both to W.W.S., SFB 1160 project C01and Deutsches Zentrum fuer Luft- und Raumfahrt, Germany (DLR, grant number 01KI2077) to M.S.

## Author contributions

I.G. performed the Jurkat experiments, and the ones with the Ramos cells, O.S.Y. and I.G. cloned the TRuC plasmids, K.C. and I.G. performed all experiments including SARS-CoV-2 whereas K.C. performed all titer analysis and N staining experiments, S.C.P. did the co-IP, P. S. and N.G. helped in the flow cytometry, D.S. performed the microscopy killing experiment, N.V. cloned the S-mScarlet fusion and generated all the S-expressing Ramos cells, J.S. cloned the Luciferase-BFP plasmid, M.R., M.S. and W.W.S. supervised the work. All authors analysed data and helped in preparing the manuscript.

## Additional information

Correspondence and requests for materials should be addressed to W.W.S.

## SUPPLEMENT

### Content

Figures S1, S2, S3, S4, S5, S6 and S7

**Fig. S1.**
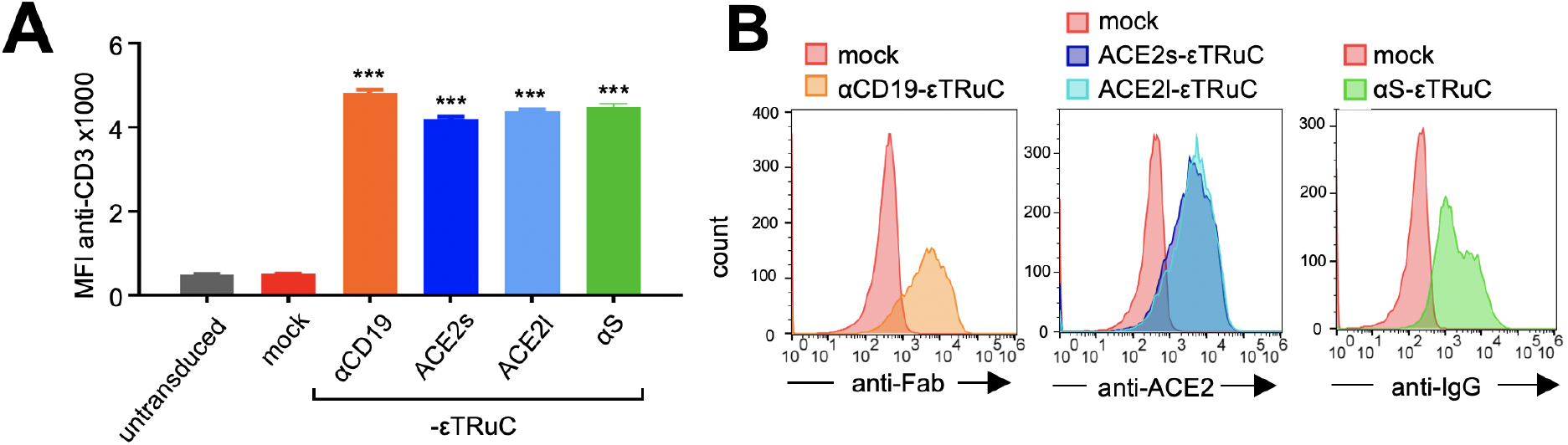
The S-specific ε TRuCs are expressed on Jurkat T cells. **A**, The Jurkat CD3εKO transductants as indicated were stained with an anti-CD3 antibody and measured by flow cytometry. The MFI of the CD3 stain is shown. This is a quantification of triplicates, as shown in figure 1C. **B**, The cells from A were stained with an anti-ACE2, anti-IgG, or anti-Fab antibody and measured by flow cytometry (the experiment was performed more than three times, n>3). These are some of the original data that were used to generate figures 1D and 1E. These data show that the αCD19-, ACE2s-, ACE2l- and the αS-εTRuC are expressed on the surface of Jurkat CD3εKO cells.

**Fig. S2.**
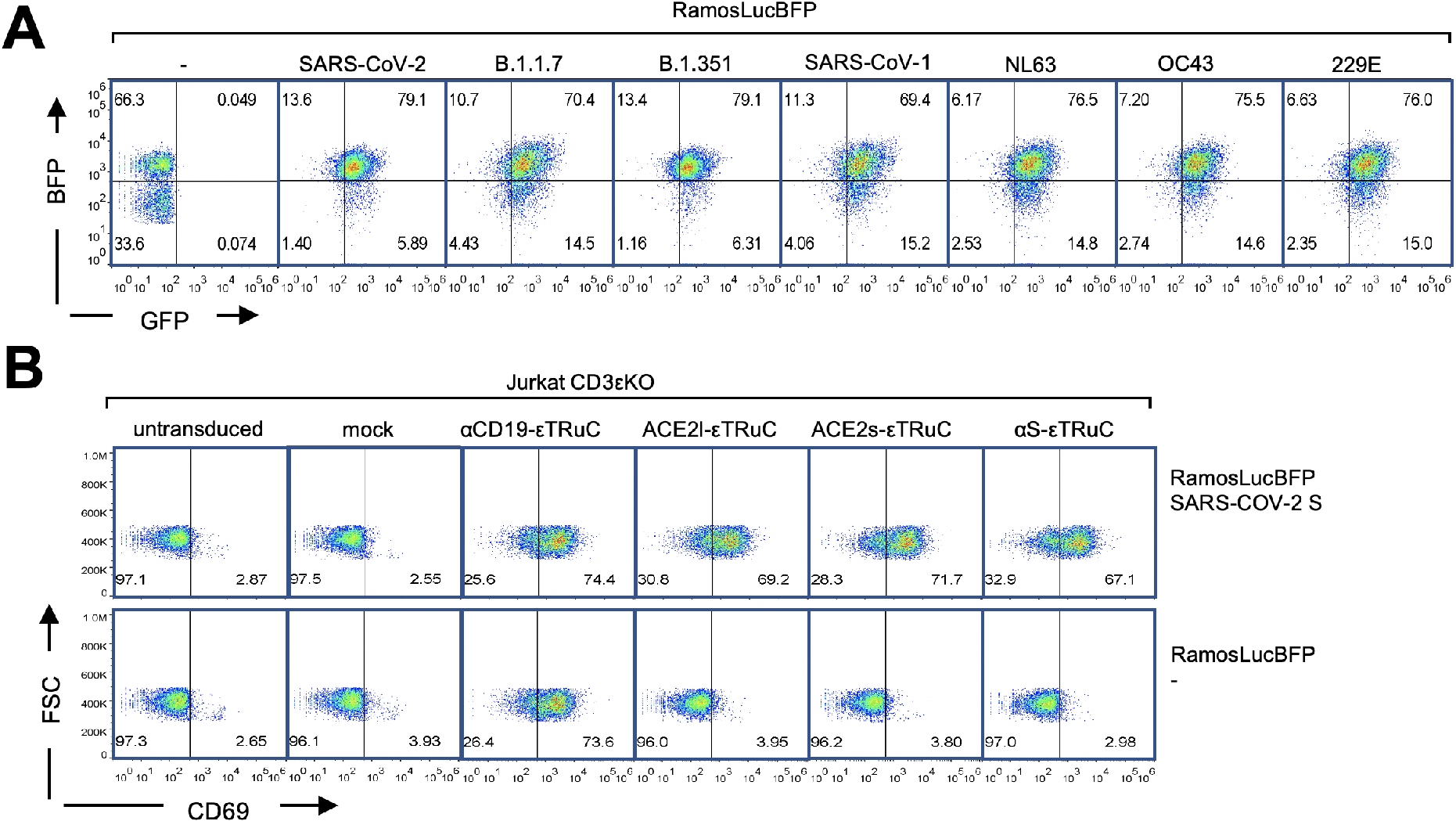
The S-specific ε TRuC T cells bind to the S and are activated by S-expressing Ramos cells. **A**, Ramos cells expressing the S of the corona viruses as indicated and GFP were lentivirally transduced to bicistronically express firefly luciferase and BFP. The flow cytometric plots show the GFP and BFP fluorescence intensity (the experiment was performed more than three times, n>3). This data demonstrates that the Ramos cells express both the S and luciferase. **B**, Ramos cells without S and those expressing SARS-CoV-2 S were co-cultured for 9 h with the Jurkat CD3εKO cells expressing the εTRuCs as indicated, being transduced with the GFP-expressing mock vector or left untransduced. Subsequently, cells were stained with anti-CD69 and anti-CD3 antibodies and analysed by flow cytometry. These original data were used to generate figure 2C.

**Fig. S3.**
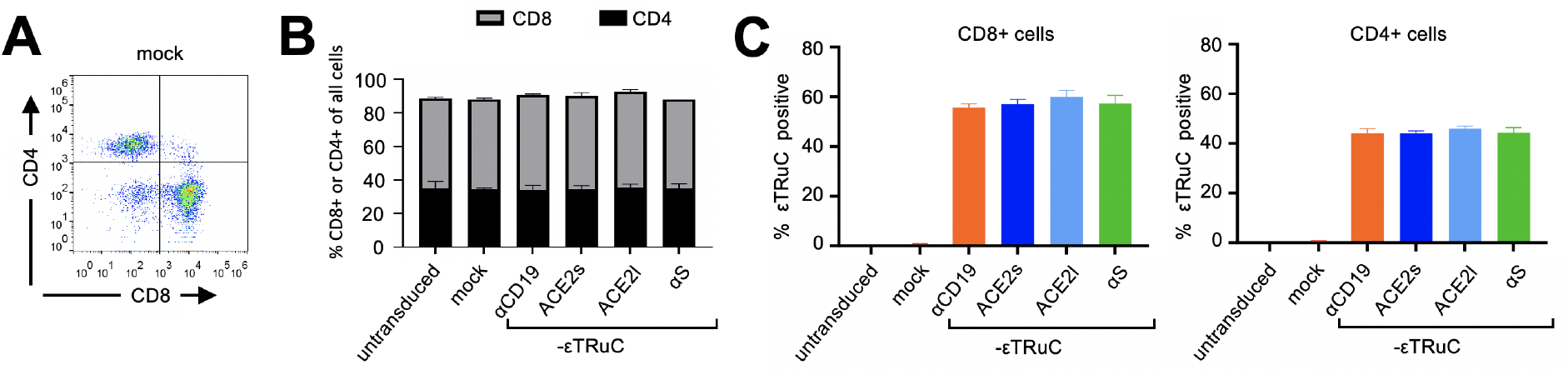
The S-specific ε TRuCs are expressed on CD4^+^ and CD8^+^ T cells. **A**, At day 5 after mock or εTRuC transduction the primary, IL2-expanded T cells were stained with anti-CD4, anti-CD8 and the εTRuC-specific antibodies as in figure 3A. As an example, the anti-CD4 and anti-CD8 stain is shown for the mock transduction. **B,** The ratio of CD8 to CD4 cells, of the untransduced or transduced primary T cells is shown from the data as in A. **C,** The percentage of CD8^+^TRuC^+^ (left panel) and CD4^+^ TRuC^+^ (right panel) cells are shown from the data as in A (n>3). These data show that the εTRuC expression did not alter the ratio of CD8^+^ to CD4^+^ T cells.

**Fig. S4.**
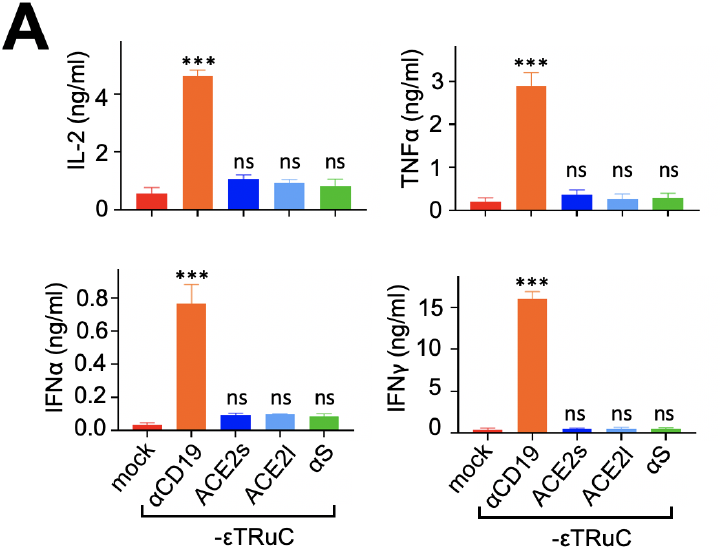
The S-specific ε TRuC T cells do not secrete cytokines upon stimulation with Ramos cells that do not express any S. **A**, Secretion of cytokines by the εTRuC T cells following co-culture with the Ramos cells not expressing SARS-CoV-2 S was quantified by ELISA in triplicates (n=3). This was done in parallel to data shown in figure 4B and demonstrates that the engineered T cells are specifically activated by cells expressing S.

**Fig. S5.**
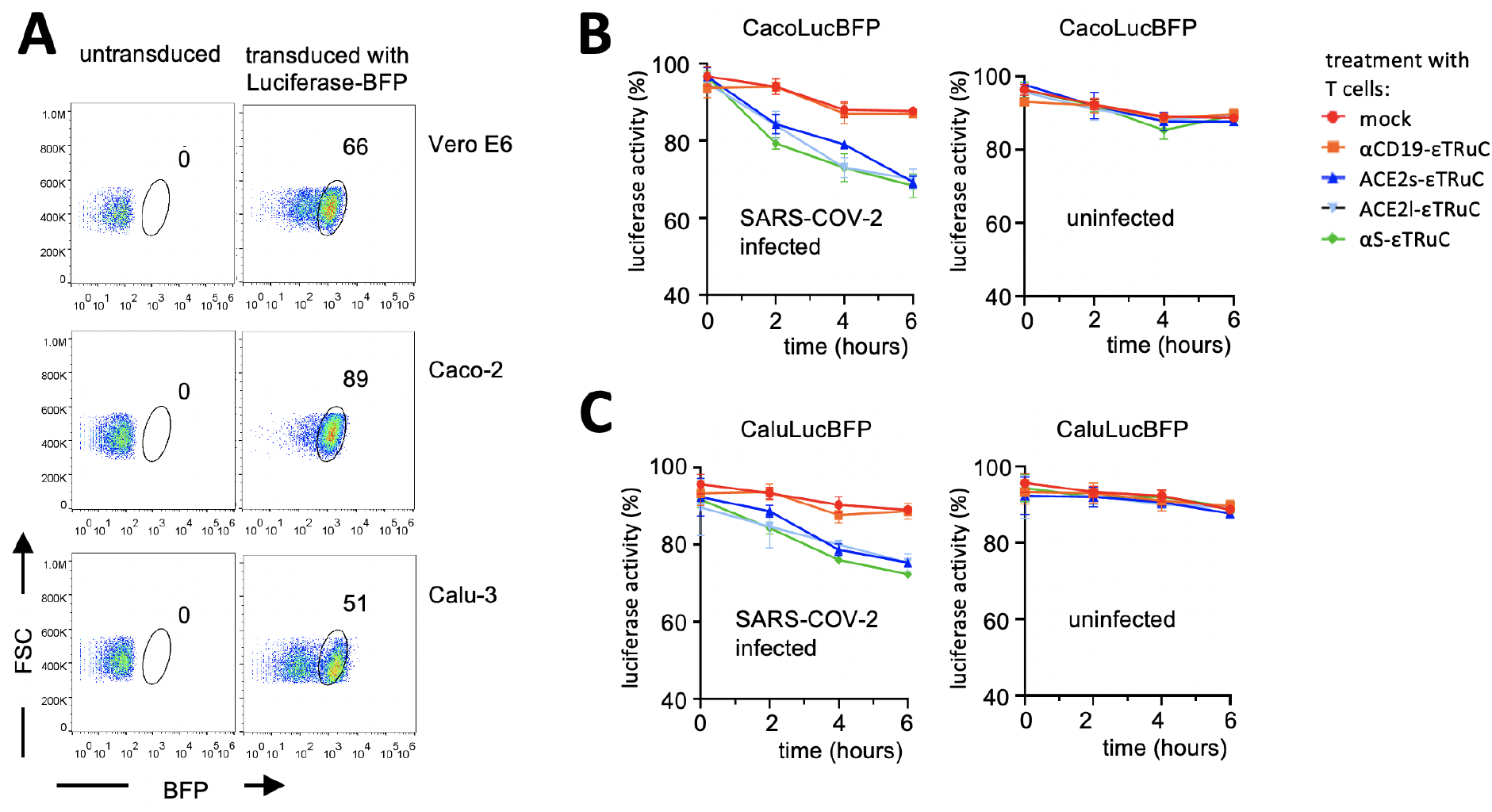
Infected CacoLucBFP and CaluLucBFP cells are killed by S-specific ε TRuC T cells. **A**, Vero E6, Caco-2 and Calu-3 cell lines were transduced to express luciferase-BFP or left untransduced and analysed by flow cytometry (n=7). In fact, 50-90% of the resulting VeroLucBFP, CacoLucBFP and CaluLucBFP cells express luciferase-BFP. These cells were used for all the SARS-CoV-2 infection experiments. **B**, The left panel shows the luciferase activity of CacoLucBFP cells when infected with SARS-CoV-2 (MOI 0.05) and treated with the primary εTRuC T cells as indicated. The right panel shows the same treatment but using uninfected CacoLucBFP cells (n=2). **C**, Data shown are from an identical experiment as in (B), but with CaluLucBFP instead of the CacoLucBFP cells (n=2). This shows that SARS-CoV-2-infected Caco-2 and Calu-3 cells are killed by the primary S-specific εTRuC T cells.

**Fig. S6.**
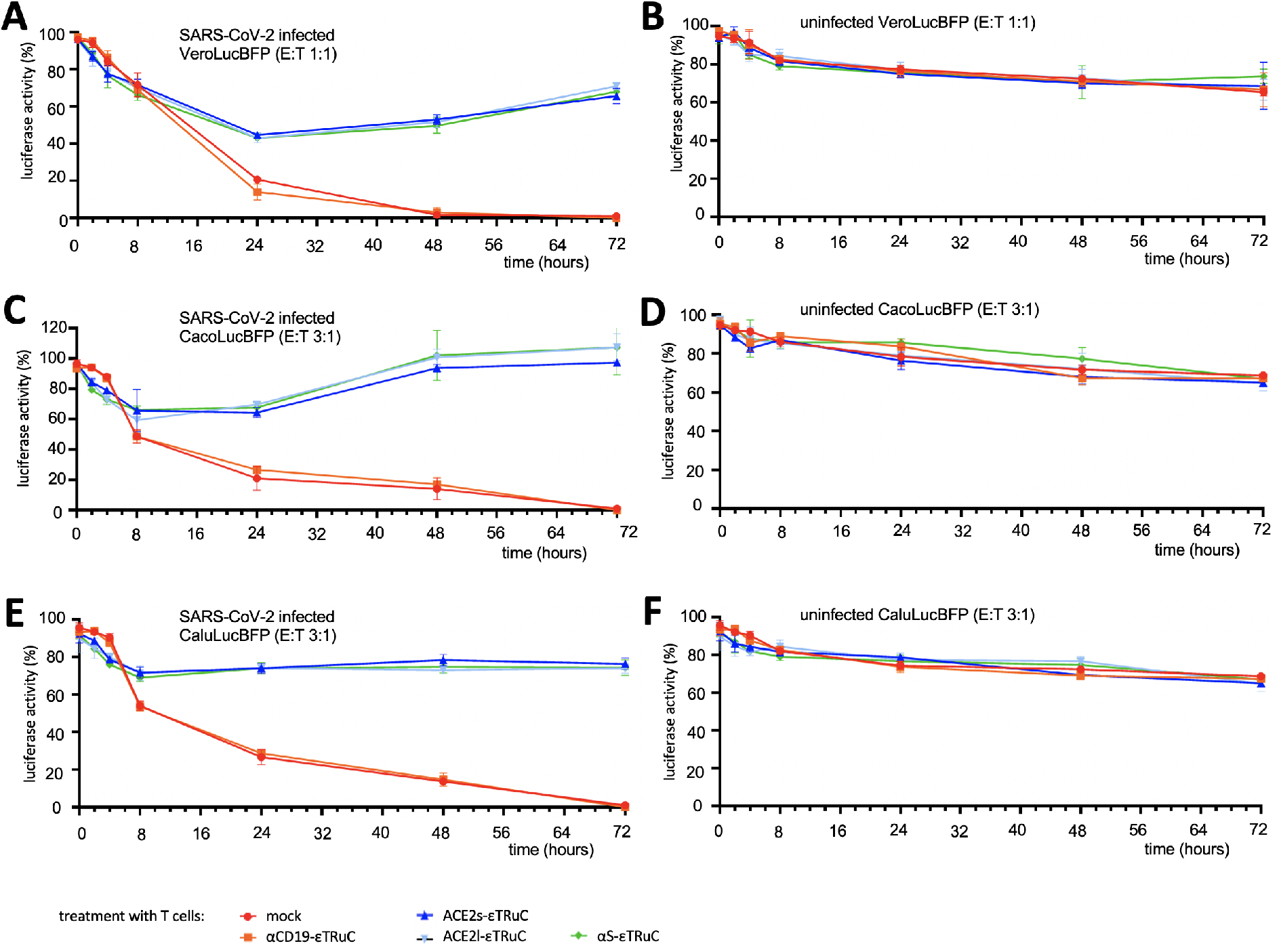
Prevention of luciferase loss in infected Caco-2 or Calu-3 cells by the εTRuC T cell treatment. **A,** VeroLucBFP cells expressing luciferase were infected with MOI 0.05 of SARS-CoV-2 virus and treated with mock transduced, αCD19-εTRuC, ACE2s-εTRuC, ACE2l-εTRuC, and αS-εTRuC expressing primary T cells in the ratio of 1:1, i.e., one transduced T cell was added per VeroE6 cell. **B,** The negative control is the treatment of Vero E6 cells without infection but treatment with mock, αCD19-εTRuC, ACE2s-εTRuC, ACE2l-εTRuC, and αS-εTRuC primary T cells in 1:1 ratio. **C,** CacoLucBFP cells infected with SARS-CoV-2 virus (MOI 0.05) were treated with mock, αCD19-εTRuC, ACE2s-εTRuC, ACE2l-εTRuC, and αS-εTRuC primary cells in the ratio of 3:1, i.e., three transduced T cells were added per target cell. **D,** The negative control for is the treatment of CacoLucBFP cells without infection but treatment with mock, αCD19-εTRuC, ACE2s-εTRuC, ACE2l-εTRuC, and αS-εTRuC primary T cells in 3:1 ratio. **E,** CaluLucBFP cells infected with SARS-CoV-2 virus (MOI 0.05) were treated with mock, αCD19-εTRuC, ACE2s-εTRuC, ACE2l-εTRuC, and αS-εTRuC primary cells in the ratio of 3:1, i.e., three transduced T cells were added per target cell. **F,** The negative control for Fig S6E is the treatment of Calu-3 cells without infection but treatment with mock, αCD19-εTRuC, ACE2s-εTRuC, ACE2l-εTRuC, and αS-εTRuC primary cells in 3:1 ratio.

**Fig. S7.**
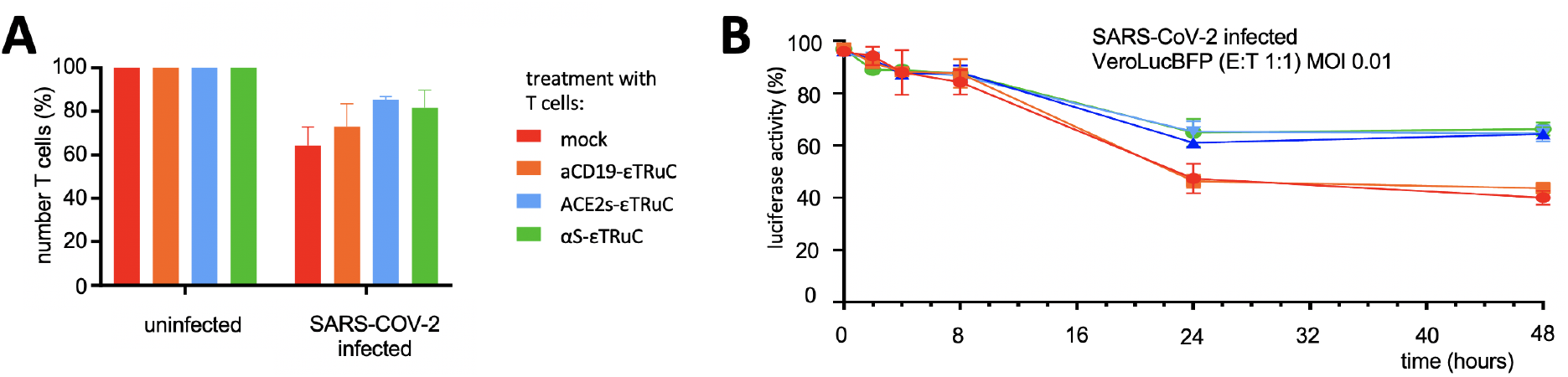
SARS-CoV-2 infection reduces the number of VeroLucBFP cells at 72 h post infection independent of the T cell treatment. **A,** The number of infected VeroLucBFP cells measured by flow cytometry along with the data of figure 7B are shown as a relative value to the uninfected VeroLucBFP cells measured after 72 h of infection. The data show that the number of cells was unaffected by our εTRuC T cell treatment. **B,** The luciferase activity of VeroLucBFP cells infected with SARS-CoV-2 (MOI 0.01) and treated with the εTRuC primary T cells at ratio of T cell : VeroLucBFP cell of 1:1 (E:T) is shown. The data demonstrate that at low MOI the initial killing of infected cells cannot be detected, but the prevention of a partial translational shutdown took place.

